# Polarity-dependent modulation of sleep oscillations and cortical excitability in aging

**DOI:** 10.1101/2025.07.03.662520

**Authors:** Buse Dikici, Robert Malinowski, Jan-Bernhard Kordaß, Klaus Obermayer, Julia Ladenbauer, Agnes Flöel

**Affiliations:** Department of Neurology, Universitätsmedizin Greifswald, Greifswald, Germany; Technische Universität Berlin, Berlin, Germany; Bernstein Center for Computational Neuroscience Berlin, Berlin, Germany

**Author notes:** shared last authorship (Julia Ladenbauer, Agnes Flöel). Corresponding authors. (J. Ladenbauer), (A. Flöel). Correspondence Agnes Flöel, Universitätsmedizin Greifswald, Klinik und Poliklinik für Neurologie, Ferdinand-Sauerbruch-Straße, 17475 Greifswald, Germany.

## Abstract

During non-rapid eye movement (NREM) sleep, cortical slow oscillation (SO; <1 Hz) and thalamic sleep spindle activity (12-15 Hz) interact through precise phase coupling to support memory consolidation. Slow oscillatory transcranial direct current stimulation (so-tDCS) can modulate these oscillations. Traditionally, anodal so-tDCS is used to depolarize the cortex during SO up-states, thereby promoting SO activity and SO-spindle coupling. However, intracranial findings suggest that SO down-states, characterized by cortical hyperpolarization, can trigger thalamic spindle bursts. This raises the hypothesis that cathodal so-tDCS, by promoting hyperpolarization, could selectively enhance down-states and more effectively improve SO-spindle coupling.

We tested this hypothesis in twenty-two healthy older adults, a population known to exhibit diminished NREM oscillatory activity. Each participant received cathodal, anodal, and sham so-tDCS in separate nap sleep sessions. We quantified SO and spindle characteristics, their temporal coupling, and cortical excitation/inhibition (E/I) balance using EEG spectral slope. We also assessed individual circadian preference (chronotype) as a potential moderator.

We found that anodal so-tDCS improved SO-spindle synchrony, and increased spindle power over sham in participants with intermediate or evening chronotypes, while cathodal so-tDCS did not enhance these oscillatory measures compared to sham, despite prolonging SO down-states. Anodal stimulation also elevated E/I balance, indicating increased cortical excitability, whereas cathodal stimulation did not produce the anticipated opposite shift. In summary, anodal, but not cathodal so-tDCS, effectively enhanced thalamocortical interactions underlying memory consolidation. Furthermore, these findings highlight the importance of individual factors such as chronotype in brain stimulation responsiveness.

## Introduction

Cortical slow oscillations (SOs; <1 Hz) and thalamocortical sleep spindles (12-15 Hz) are core features of non-rapid eye movement (NREM) sleep. These rhythms are not only temporally structured but also functionally interdependent: SOs temporally coordinate the emergence of spindles, with spindle bursts preferentially occurring near the SO up-state, when cortical excitability is high ^1^. This precise phase coupling is thought to underlie systems-level memory consolidation, and was shown to be diminished in older adults ^2,3^. The ability to modulate these oscillatory dynamics noninvasively has opened new avenues for probing and potentially enhancing sleep-related memory consolidation. A promising non-invasive approach is slow oscillatory transcranial direct current stimulation (so-tDCS) with anodal polarity ^4,5^. Delivered bifrontally during NREM sleep, the sinusoidal current with SO frequency (< 1 Hz) is thought to depolarize cortical neurons towards firing threshold ^6^, thereby facilitating SO up-states and entraining the brain’s endogenous SO rhythm ^7^. Consistent with this mechanism, anodal so-tDCS has been demonstrated to enhance SO activity while also promoting spindle activity ^7–10^ and improving their precisely timed phase coupling ^11,12^. Concurrently, anodal so-tDCS has been shown to improve memory retention performance, with SO-spindle coupling being most informative for this improvement ^5^.

The rationale for using anodal polarity was originally grounded in observations of surface negative direct current (DC) potential shifts during transitions into slow wave sleep ^13^. These shifts suggest increased cortical excitability and showed strong associations with spindle and slow wave activity during NREM sleep ^14^. As a result, anodal so-tDCS has become the standard tCS approach for enhancing NREM sleep oscillations during sleep (e.g., ^8,9,11^).

However, recent findings from intracranial recordings in humans challenge the choice of anodal polarity for so-tDCS application. Mak-McCully and colleagues ^15^ identified cortical hyperpolarized down-states during SO events as key triggers for thalamic down-states, which in turn initiate spindle bursts that feed back to the cortex, timing cortical spindle activity to the late rising phase of the SO. These results suggest that the occurrence and timing of thalamo-cortical spindles is strongly affected by cortical hyperpolarizations that produce thalamic down-states. From this mechanistic perspective, biasing cortical networks toward hyperpolarization during the down-state via cathodal so-tDCS could theoretically engage this cascade more directly than promoting cortical depolarization.

Moreover, stimulation approaches that enhance cortical hyperpolarization may be especially beneficial in aging populations. Neuronal “hyperactivity” (i. e., more cortical depolarization) is commonly observed in older compared to younger adults ^16^, and is especially pronounced in pathological aging, such as Alzheimer’s disease ^17^. This hyperactivity reflects a disruption in cortical excitation/inhibition (E/I) balance, a process associated with cognitive decline ^18^. By promoting hyperpolarizing phases, cathodal so-tDCS may further help restore E/I balance and potentially reduce age-related cognitive impairments ^19^.

In the present study, we investigated the effects of cathodal so-tDCS on NREM sleep oscillations in a within-subject experiment in healthy older adults. Our goals were three-fold: first, to determine the influence of cathodal so-tDCS during a 90-minute daytime nap on (i) SO down-states, (ii) spindle activity, (iii) precision of slow oscillation-spindle coupling, and (iv) cortical E/I balance indexed by EEG spectral slope as an indication for changes in cortical excitability ^20^; second, to compare effects of cathodal and anodal so-tDCS (both relative to sham) on sleep microstructure; and third, given recent evidence that individual circadian preference (chronotype) can influence neuroplastic responses to continuous tDCS during wakefulness ^21^, to assess whether individual circadian preference moderates the impact of so-tDCS effects of either polarity during sleep.

## Materials and Methods

### Participants

Twenty-seven healthy older adults (age 55-79, mean 65.9 ± SD 6.8 years) participated in the cathodal so-tDCS condition of a larger study examining four tES protocols. Inclusion required sufficient sleep during previous nap sessions (with anodal so-tDCS and sham), defined as completing at least seven so-tDCS /sham blocks. Six participants were excluded due to insufficient sleep during the cathodal so-tDCS session, leaving 22 participants for final analysis. Participants were screened for eligibility via structured telephone interview to ensure that they did not have a history of severe untreated neurological, psychiatric, and sleep disorders. On site, exclusion criteria additionally encompassed intake of medications acting on the central nervous system (e.g., antipsychotics, antidepressants, antihistamines, benzodiazepines, or any sleep-inducing medications), severe hearing or vision impairment, alcohol or substance abuse, a contraindication or inability to undergo MR imaging, and brain pathologies identified on MRI scan (brain tumour and stroke), as well as a lack of German language skills. Furthermore, participants underwent extensive baseline assessments (see Supplemental Information for more details and Table S1 for baseline characteristics). The study was ethically approved, adhered to the Declaration of Helsinki, and all participants provided written informed consent and received compensation.

### Procedure

Nap sessions took place at the sleep laboratory of the Department of Neurology, University Medicine Greifswald, Germany. Following an adaptation nap, participants received three different stimulation conditions in a randomized order: anodal so-tDCS, anodal tDCS, and sham condition, followed by the cathodal so-tDCS condition. Each session was separated by at least one week.

Upon arrival at the sleep laboratory at 11:30 a.m., participants were prepared for EEG recordings and then completed the learning phase of two computerized declarative memory tasks consisting of an object-location task and a verbal paired-associate learning task. At 2:00 p.m. participants were asked to attempt to sleep for a period of 90 minutes, and subsequently, participants performed the retrieval phase of the memory tasks. The last cathodal so-tDCS session was conducted in an identical manner with the exception that the experimental session began at 12:30 p.m., as no memory tasks were tested for this condition. Please see Supplemental Information for more details on procedure and analyses.

### Brain Stimulation

A setup of two battery-driven stimulators (DC-Stimulator; NeuroConn, Ilmenau, Germany) were used to deliver the oscillatory current stimulation. Stimulation electrodes (8 mm diameter) were placed bilaterally at frontal locations (F3, F4), with return electrodes on ipsilateral mastoids (8 mm diameter; M1, M2). The current, of either anodal or cathodal polarity followed a sinusoidal waveform oscillating between 5 and 255 µA, and between -5 and -255 µA, respectively, generating a maximum current density of 0.507 mA/cm² per hemisphere. Stimulation began four minutes after stable NREM stage 2 (N2) sleep, delivered in repeated 1-minute blocks (7-15 blocks, min. 7 were required for inclusion), each separated by at least 1.5 minutes. Stimulation was suspended if participants transitioned out of N2/N3, and it was resumed after stable return. Sham sessions followed identical temporal patterns without actual stimulation. After all naps, participants were asked whether they had perceived any sensations during the naps.

### Sleep monitoring and EEG preprocessing

The EEG cap (actiCAP, Brain Products GmbH, Gilching, Germany; 64-channels) for sleep monitoring was prepared according to the 10-20 international EEG system after tracking the electrode positions on the cap with a neuronavigation system (Rogue Research, Montréal, Canada) to ensure identical cap and stimulation electrode placement in each nap session. During recording EEG data were referenced to the nose electrode, with EOG and chin EMG recorded for sleep stagin (500 Hz sampling rate). After recording, the YASA sleep scoring algorithm ^22^ was used to conduct offline sleep staging on 30 second epochs, which was verified by an expert scorer. As AASM recommended EEG derivations were partly occupied by stimulation electrodes, we used the following channels for offline sleep staging: F1, C3, O1, or F2, C4, O2 in case of noisy channels; in addition to EOG and MEG. Stimulation epochs were not scored due to strong artefacts in the EEG signal, a method also applied to the corresponding epochs in the sham session.

Preprocessing and analysis of EEG data was conducted with MNE-python (v1.7.1) ^23^. After applying a notch filter at 50 Hz and its harmonics, data was downsampled to 200 Hz, followed by automated annotation of so-tDCS/sham intervals, bad-channel identification (ANOAR package ^24^), and visual artifact rejection.

### EEG analyses

Analyses focused primarily on 1-minute intervals following each so-tDCS or sham block at fronto-central derivations (Fz, or FC1 or FC2 in case of noise).

#### Spectral power analyses

YASA ^22^ was used to perform spectral power analysis in the SO (0.5-1Hz) and spindle frequency range (12-15 Hz). Power spectral density (PSD) was computed using Welch’s method on consecutive 4-second epochs with 50% overlap and a Hamming window, resulting in a frequency bin resolution of 0.25 Hz. The absolute band power for SO (0.5-1 Hz) and spindle (12-15 Hz) frequency ranges were calculated by integrating the power spectrum within these frequency bands (area under the curve). To obtain relative power, the absolute band powers were then normalized by dividing by the total power within the 0.5-35 Hz frequency range.

#### SO event detection

SO detection followed practices established in the literature ^5,12,25^. In brief, the signal was filtered using a finite impulse response (FIR) bandpass filter with a frequency range of 0.16-1.25 Hz, after which all negative-to-positive zero-crossings in the signal were identified. The time between two successive negative zero-crossings was measured, as well as the amplitude difference between the most negative point (down-state trough) after the first positive-to-negative zero-crossing, and the most positive point (up-state peak) after the next negative-to-positive crossing. SO events were marked if the amplitude difference between peak and troughs was greater than the threshold (65^th^ percentile) among all detected zero-crossings and the duration between two consecutive positive-to-negative zero-crossings was between 0.8 and 2 seconds.

#### Time-frequency representation analysis

Time-frequency representations (TFRs) were calculated for each SO event epoch (5 sec epoch) using the morlet wavelet transform with 5 cycles, after data were first down-sampled to 50 Hz. It was applied for the spindle band (10-20 Hz) in steps of 0.2 Hz to provide robust power estimates. The first and last 150 ms of each epoch were cut off to remove edge effects. After calculating TFR within the spindle band for each SO epoch, baseline adjustment followed the z-score method, such that the mean and standard deviation of the time period -2.35 to -1.5s were used in a z-score transformation of the entire SO epoch. First, a two-tailed paired-samples t-test was conducted to determine statistical significance between the conditions at the group level. Next, a cluster-based permutation test (1024 iterations) was applied using the threshold-free cluster enhancement (TFCE) method to account for multiple comparisons and to detect significant clusters. The significance threshold was set at 0.05.

#### SO-spindle coupling

In addition to TFR analyses, slow oscillation-spindle coupling was quantified through event-locked phase-amplitude coupling analyses. First, the normalized SO event segments were filtered in the SO band (0.5-1.25 Hz) and the instantaneous phase angle was extracted using a wavelet transformation. Then, the same segments were filtered in the spindle band (12-15 Hz) and the instantaneous amplitude was extracted also with a wavelet transformation. To avoid filter edge artifacts, we only considered the time range −2 to 2 s. We next extracted the maximal spindle amplitude and corresponding SO phase angle for every subject, and SO event. This yielded a distribution of spindle-at-peak phase angles for each participant and condition (e.g. 0° = positive peak/up-state, 180° = negative trough/down-state). From these distributions we computed two measures: (1) the mean coupling phase angle and (2) the coupling strength, quantified as the resultant vector length. Mean coupling phase angles and coupling strength were compared across conditions using circular statistics (Watson-Williams tests) and repeated-measures ANOVA (followed by planed contrasts (cathodal vs. cham; anodal vs. sham) in case of significant difference), respectively. Signal decomposition was carried out with the Python package Tensorpac 0.6.5 27.

#### Cortical excitation/inhibition (E/I) balance

To test whether cathodal vs. anodal so-tDCS stimulation induced divergent shifts in cortical excitability during post-stimulation intervals, we extracted the spectral slope x (the negative exponent of the 1/fx decay function) from 1-45 Hz frequency range of PSD using FOOOF package ^26^ with the ‘fixed’ aperiodic mode (similar to ^27^). The spectral slope of the log-log PSD was estimated for each interval and averaged per condition within each subject. A more negative spectral slope (i.e. a steeper decline of power at higher frequencies) indicates reduced high-frequency activity relative to low-frequency – often interpreted as a shift towards greater inhibition relative to excitation, whereas a flatter slope implies more high-frequency power and potentially greater excitatory tone. We statistically compared slopes between conditions with repeated-measures ANOVA, followed by planed contrasts (cathodal vs. cham; anodal vs. sham) in case of significant difference, and additionally checked 1min pre-stimulation baseline segments to confirm no inherent differences before stimulation. Correlations between spectral slope, spindle activity, and coupling measures were also explored, alongside potential moderation effects of individual circadian preference, which was assessed using the German version of the Morningness-eveningness questionnaire (D-MEQ) ^28^. We considered MEQ score as a continuous and as a categorical covariate (two groups), where higher scores indicated morning-type (> 58) and lower scores indicated intermediate-to evening-type. Statistical analyses were conducted using Python libraries (numpy, pandas, pycircstat2) and SPSS (Version 29.0), with significance set at p < 0.05, adjusted for multiple comparisons (Holm-Bonferroni).

## Results

### Cathodal so-tDCS effects

We first focused on the effects of cathodal so-tDCS on SO parameter. Cathodal so-tDCS did not influence the primary SO amplitude measure in this evaluation: the magnitude of SO down-states (F_(2, 42)_ = 0.961, p = 0.37, Greenhouse-Geisser corrected). Further, no significant changes were observed in SO density, SO up-state amplitudes, slope of SO events, or the overall peak-to-peak SO amplitudes relative to sham (all p-values > 0.1). Similarly, spectral SO power remained unaffected by cathodal so-tDCS (F_(2, 42)_ = 1.419, p = 0.25). However, cathodal so-tDCS strongly affected the duration of negative SO periods, resulting in prolonged negative SO durations (F_(2, 42)_ = 6.132, p = 0.005, Holm-Bonferroni adjusted p-value: cathodal-sham: p-adj = 0.015, Cohen’s d = 0.668, Fig. 1a), indicating a prolonged hyperpolarizing phase of the SO cycle. In addition, positive SO duration was shortened after cathodal so-tDCS relative to sham (F_(2, 42)_ = 6.432, p = 0.004; anodal-sham p-adj = 0.032, Cohen’s d = 0.56, Fig.1 b).

**Figure 1.**
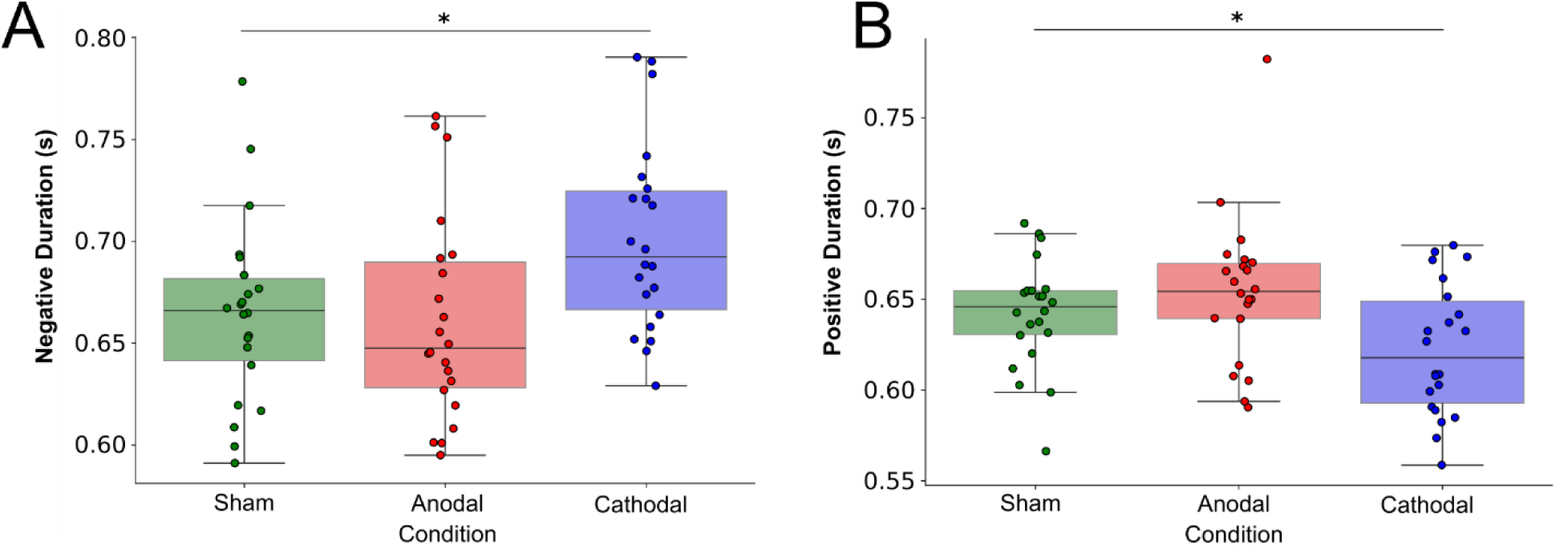
Cathodal so-tDCS modulates SO durations. A. Negative and B. positive SO duration by condition. Cathodal so-tDCS prolonged negative SO durations and shortened positive SO durations relative to sham. *p < 0.05

In terms of sleep spindle activity, cathodal so-tDCS did not significantly alter spindle power relative to sham (F_(2, 42)_ = 1.554, p = 0.223). Furthermore, no significant effects of cathodal so-tDCS were observed on SO-spindle coupling. Specifically, it did not increase spindle activity during the SO upstate in the TFR analysis relative to sham (see Fig. 2), and circular analyses of mean coupling direction of spindles within the SO cycle showed only minimal differences compared to sham (circular Watson-Williams test: p = 0.794; Cohen’s d = 0.1208, mean phase difference = 2.21° [95% CI: -13.47° to 17.89°]), indicating a negligible overall cathodal so-tDCS effect on temporal synchrony between SO and spindle activity. Consistent with these findings, cathodal so-tDCS showed no impact on SO-spindle coupling strength (F_(2,42)_ = 0.020, p = 0.98).

**Figure 2:**
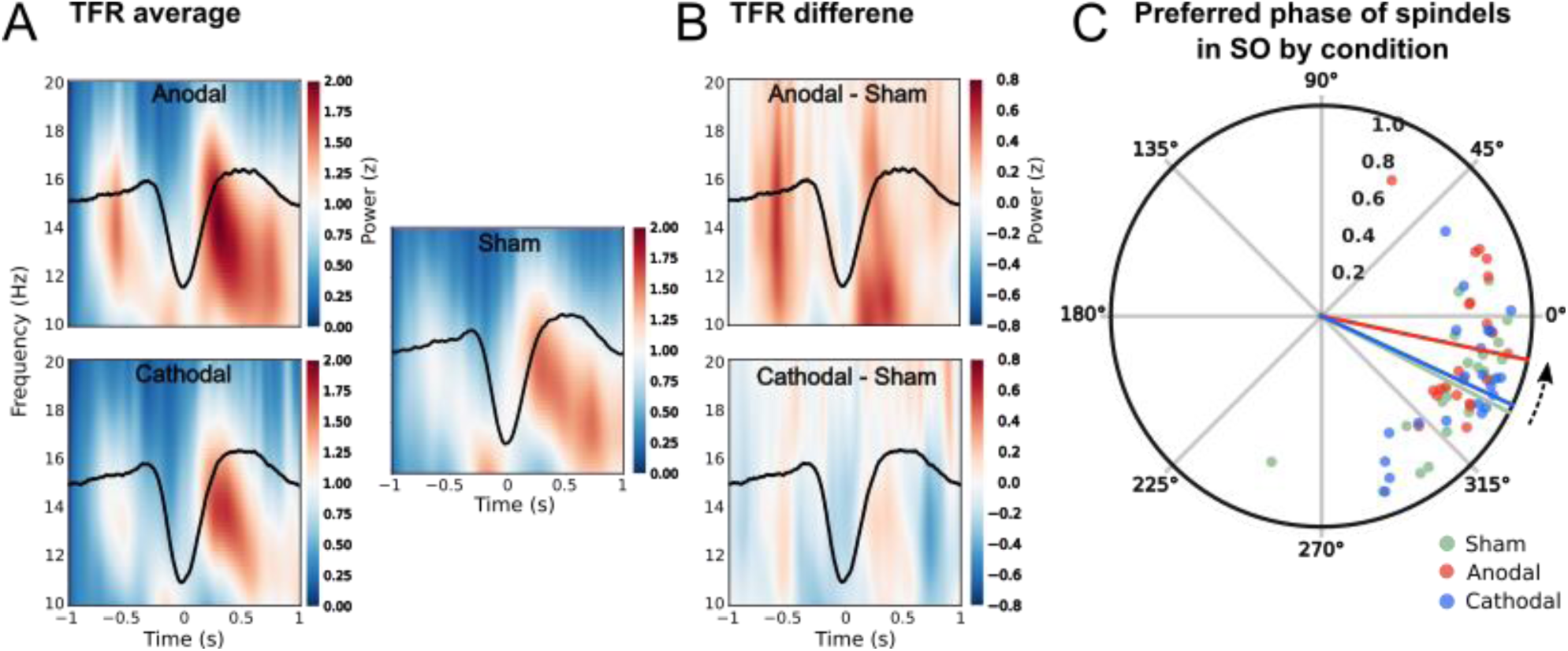
Stimulation polarity impacts on slow oscillation (SO)-spindle coupling. A. Time-frequency representations (TFRs) of SO-locked power, averaged across subjects for each stimulation condition, shown as change from pre-event baseline. B. Statistical map of condition differences in SO-locked power. Note that spindle-band (12-18 Hz) activity during SO up-phases increases in the anodal condition, but not in the cathodal condition, relative to sham. C. Polar plot showing the mean SO phase at which spindle power peaks for each condition (colored lines), with individual subject averages represented by dots for each condition: sham (green), anodal (red), and cathodal (blue). Note that spindle timing in the anodal so-tDCS condition is shifted toward the SO up-state (0°), indicating enhanced phase synchrony compared to sham and cathodal so-tDCS.

To assess effects on cortical excitability, the slope of the EEG power spectral density was used as a proxy measure of the E/I balance ^20^. Although a slight increase in spectral slope was observed under cathodal so-tDCS relative to sham, this change failed to reach statistical significance (F_(2, 42)_ = 6.121, p = 0.005, partial η² = = 0.226; planed contrast cathodal vs. sham: p-adj = 0.073; Cohen’s d = 0.402).

### Anodal so-tDSC effects

Anodal stimulation demonstrated a markedly different pattern of effects across the same sleep parameters. Parallel analyses for anodal so-tDCS revealed similarly negligible effects on SO parameters. Specifically, no significant alterations were observed in the magnitude of SO down-states, SO up-states, number of SO events, SO slope, peak-to-peak amplitude, or duration of positive and negative duration of the SO events compared to sham (all p > 0.1). Likewise, SO and spindle power remained unaffected by anodal so-tDCS (F_(2,42)_ = 1.419, p = 0.25; and F_(2, 42)_ = 1.554, p = 0.223, respectively). However, anodal so-tDCS facilitated SO-spindle coupling. Time-frequency representations indicated enhanced spindle power during the late rising phase of the SO following anodal so-tDCS relative to sham (see Fig. 2 a,b). Although the SO-epoch interval did not reach statistical significance after cluster-based permutation test to account for multiple comparisons, the trend clearly indicates an increase in spindle activity during the depolarizing SO up-phase. Complementary circular statistical analyses confirmed a strong shift in the preferred spindle phase after anodal so-tDCS compared to sham, with spindles occurring closer to the SO upstate peak (mean angle ± SD: anodal = 348.0° ± 26.25°; sham = 332.84° ± 27.24°), indicating improved precision of spindle alignment relative to sham (Fig. 2c). The mean difference (15.16°; 95% CI: -0.65° to 30.97°) indicated a large effect size (Cohen’s d = 0.824), though the difference marginally missed statistical significance (Watson-Williams test: p = 0.081). Coupling strength remained unchanged (F_(2, 42)_ = 0.020, p = 0.98), suggesting that the effect of anodal so-tDCS relative to sham was specific to timing, rather than general coupling strength.

Further significant results emerged in the context of cortical excitability, where anodal so-tDCS produced a robust increase in E/I balance relative to sham (F_(2,42)_ = 6.121, p = 0.005; p-adj = 0.004, Cohen’s d = 0.752, see Fig. 3), indicating enhanced cortical excitability. Baseline spectral slopes prior to stimulation onset did not differ between conditions (F_(2, 42)_ = 0.226, p = 0.734).

**Figure 3:**
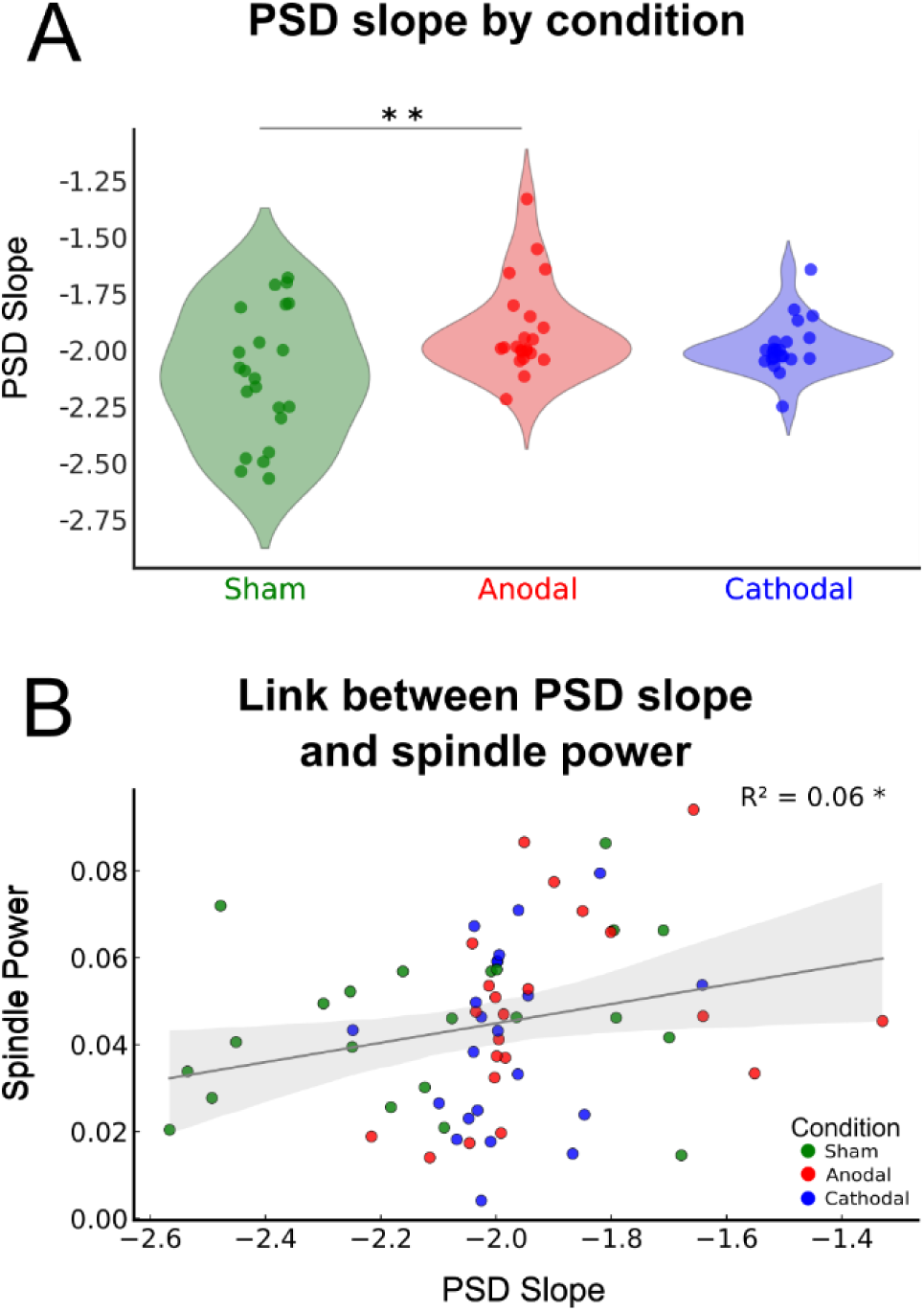
So-tDCS effects on excitation/in-hibition (E/I) balance and PSD slope associ-ation with spindle power. A. Power spectral density (PSD) slope across conditions. Note that PSD slope is increased (less negative) after anodal so-tCS relative to sham. Cathodal so-tDCS shows a similar trend. B. Positive association between PSD slope and spindle power across conditions, suggesting that E/I balance contributes to spindle power expression, though with very modest explanatory variance. *p < 0.05, **p < 0.01

A comparative evaluation highlighted polarity-specific effects of so-tDCS on NREM sleep microstructure. While only cathodal so-tDCS influenced SO durations, with prolonged negative SO phases and shortened positive SO phases, anodal so-tDCS markedly promoted SO-spindle coupling, evident by increased spindle activity during the SO up-state as well as improved synchrony of SO-spindle coupling. In addition, anodal so-tDCS markedly shifted E/I balance in the direction of increased excitability. In contrast, cathodal so-tDCS failed to produce significant effects on these outcomes, although there was a trend in the same direction especially for E/I balance.

### Morning-evening preference as a moderator

Chronotype, specifically the continuous degree of preference for morning or evening activity (i.e. MEQ score), emerged as an important moderator of anodal so-tDCS effects on spindle activity. We observed an interaction effect between stimulation condition and morning-evening preference (i.e. MEQ score; F_(2, 40)_ = 4.066, p = 0.038, Greenhouse-Geisser corrected, partial η² = 0.169). Individuals with low MEQ score, classified as intermediate- to evening-types (MEQ score < 59; n = 10), showed increases in spindle power after anodal so-tDCS compared to sham (mean change ± SD: 24.1% ± 72.1%, p = 0.077). Conversely, morning-types (n = 12) experienced no spindle power benefit or decrease following anodal so-tDCS (mean change ± SD: -10.6% ± 54.5%, p = 0.246). By comparison in cathodal condition, neither intermediate- to evening-nor morning-types showed effects on spindle power compared to sham (mean change ± SD: -6.01% ± 57.88%, p = 0.575; and -11.32% ± 60.70%, p= 0.250, respectively). This finding was substantiated by a strong negative correlation between MEQ scores and individual anodal so-tDCS-induced spindle increase (r = -0.59, p = 0.004), with MEQ scores explaining 35.1% of the variance in spindle power change after anodal so-tDCS compared to sham (see Fig. 4a). For cathodal so-tDCS, the negative correlation with MEQ score demonstrated a smaller effect size that approached but did not reach statistical significance (r = -0.41, p = 0.057, R² = 0.169), indicating a similar directional relationship but with much weaker and less consistent chronotype modulation effect compared to anodal so-tDCS. No other interaction effect with chronotype was evident for all other investigated NREM sleep parameter (all p > 0.3).

**Figure 4.**
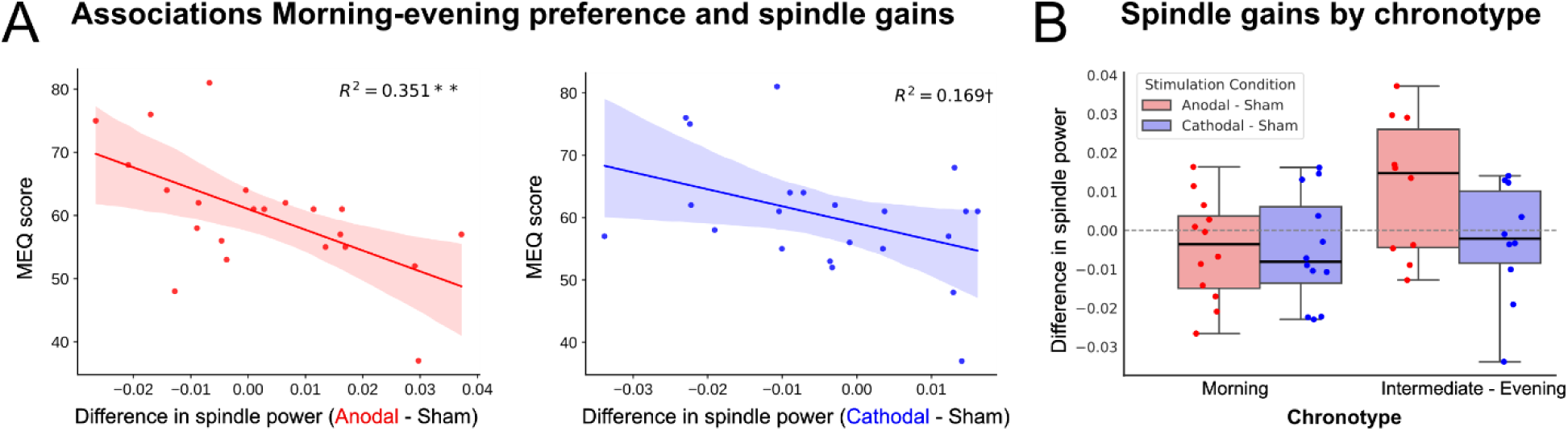
Chronotype moderates the effects of so-tDCS on spindle activity. A. Spindle gains in anodal (left) and cathodal so-tDCS (right) condition plotted against individual MEQ scores. Spindle power changes after anodal so-tDCS is strongest for intermediate- and evening preference (MEQ score < 59) relative to sham. A similar, though much weaker, trend is observed following cathodal so-tDCS. B. Spindle power changes after anodal (red) and cathodal (blue) so-tDCS (relative to sham) displayed by chronotype group. **p < 0.01. †p < 0.1

Additional regression analyses investigating the relationship between cortical excitability and sleep oscillations across conditions revealed that E/I balance significantly predicted spindle power (B = 0.022, β = 0.25, SE = 0.11, t = 2.064, p = 0.043, Fig. 3b), though with very modest explanatory variance (6.2%), while it showed no relationship with coupling phase (p = 0.289, R² = 0.018).

In addition to our primary objective of examining the impact of so-tDCS on sleep microstructure, we evaluated whether so-tDCS conditions influenced macro-sleep architecture, including sleep-stage proportions (across the entire nap and within 1-minute post-stimulation intervals), sleep efficiency, and sleep-stage latencies. Cathodal so-tDCS had no significant effects on any of these macro-sleep parameters compared to sham stimulation (all p > 0.1; see Table S2). Conversely, during anodal so-tDCS, we observed a reduced proportion of slow-wave sleep (N3), along with an increased proportion of lighter sleep (N1), suggesting a shift toward shallower sleep under anodal so-tDCS. However, this effect may partly reflect in part pre-stimulation differences, as latencies to N2 (and consequently N3) sleep onset were longer during the anodal condition, thereby biasing subsequent sleep-stage proportions. Supporting this interpretation, analyses of 1-minute post-stimulation intervals immediately following so-tDCS/sham blocks revealed no significant differences between conditions (all p > 0.1; Table S2). Notably, a trend emerged toward fewer brief awakenings (WASO) in post-stimulation intervals following anodal so-tDCS as compared to sham (Holm-Bonferroni-corrected p = 0.062), indicating that anodal so-tDCS may exert protective effects against awakenings. Nevertheless, these results collectively suggest that anodal so-tDCS may preferentially supports N2 rather than deeper N3 sleep stages.

Furthermore, the use of a state-dependent stimulation protocol requiring N2/N3 sleep may have introduced variability in the number of stimulation and sham blocks across conditions. However, the number of blocks did not differ significantly among the three conditions (Friedman test: χ²(2) = 3.534, p = 0.171; see Table S3 for number of stimulation blocks per condition).

In sum, cathodal and anodal so-tDCS elicited distinct and polarity-specific effects on sleep-related neural oscillations. While cathodal so-tDSC influenced SO durations, it did not significantly affect other sleep parameter. In contrast, anodal so-tDCS promoted more precise SO-spindle phase alignment, increased cortical excitability and enhanced spindle power in individuals with an intermediate- to evening-chronotype.

## Discussion

In the present study, we investigated the impact of so-tDCS polarity on the dynamics of NREM sleep oscillations that support memory consolidation in healthy older adults. We focused on how cathodal so-tDCS modulates cortical SO and thalamocortical spindle activity, and their phase coupling, compared these outcomes to anodal so-tDCS effects (both relative to sham), and determined the influence of chronotype on individual responses to both so-tDCS protocols. Our findings reveal a clear polarity-dependent difference: cathodal so-tDCS prolonged SO down-states and shortened SO up-states, suggesting enhanced cortical hyperpolarization, but did not impact on spindle activity nor on SO-spindle coupling. In contrast, anodal so-tDCS induced a marked shift in spindle timing toward the SO up-state, consistent with tighter and thus improved SO-spindle coupling, and led to a marked increase in cortical excitability compared to sham. Furthermore, individual circadian preference emerged as a key modulator: participants with a later chronotype showed the strongest anodal-induced increases in spindle activity.

### Cathodal so-tDCS prolonged down-state without spindle enhancement

Based on Mak-McCully et al. ^15^, our hypothesis was that cathodal so-tDCS, by hyperpolarizing cortical neurons ^29^ during the down-state, would increase down-states as compared to sham, and thereby influence spindle generation. We found that cathodal so-tDCS indeed prolonged duration of the SO down-state, which may reflect an augmented hyperpolarized state, that in turn may allow greater thalamic deactivation and a more pronounced rebound spindle burst, according to the mechanism proposed by Mak-McCully et al. ^15^. However, in our experimental setting, cathodal so-tDCS did not lead to changes in spindle power, nor in the precise timing of spindles relative to the SO cycle.

This failure to modulate spindle activity could be attributed to several factors. First, while evidence indicates that synchronized cortical down-states can influence thalamic rebound spindles ^15^, it is likely that an optimal timing and dynamic between down- and up-states is required for effective spindle initiation ^30–32^. Second, prolonging the down-state might not enhance this temporal pattern, possibly even disrupt it ^31^. Third, during natural NREM sleep, cortical neurons are already hyperpolarized during down-states ^33^, potentially limiting the additional impact of externally applied hyperpolarizing currents on the amplitude of the down-state. In this scenario, cathodal so-tDCS might offer little additive modulation due to a ceiling effect in membrane hyperpolarization.

It is also notable that we did not observe a decrease in the EEG spectral slope under cathodal so-tDCS. The spectral slope is increasingly recognized as a proxy for cortical excitability, with a more negative (steeper) slope reflecting stronger relative inhibition ^20^. The lack of spectral slope steepening suggests that cathodal so-tDCS did not meaningfully shift the E/I balance toward inhibition. In fact, the direction of change was toward slight flattening, indicating a trendwise increase in excitability. These findings align with other studies demonstrating that cathodal tDCS does not consistently produce hyperpolarizing effects ^34–36^. The inconsistency in results across studies may be attributed to various factors, including differences in brain states, targeted brain regions and tDCS dosage, which can modulate the effects of cathodal tDCS ^36^.

### Anodal so-tDCS improved SO-spindle coupling and increased cortical excitability

Anodal so-tDCS induced beneficial effects on NREM sleep oscillations, similar to previous studies: Spindle power was greater during late rising phase of the SO as compared to sham, largely consistent with prior work demonstrating anodal so-tDCS-related increases in spindle activity during SO events ^5,12^. Spindles were not only more prominent in power, but also more precisely timed within SO events: Under anodal stimulation, compared to sham, spindles occurred closer to the peak of the SO up-state. This phase shift reflects a tightening of SO-spindle coupling, a parameter known as critical for effective memory consolidation, and typically more temporally dispersed in older adults ^2,37^. Thus, while cathodal so-tDCS primarily affected the duration of SO components, without engaging downstream oscillatory processes, anodal so-tDCS modulated thalamocortical dynamics critical for memory consolidation. These findings demonstrate that polarity is a decisive factor for the effect of so-tDCS during sleep.

Anodal so-tDCS also enhanced E/I balance during NREM sleep, i.e., increased cortical excitability. This finding aligns well with previous reports demonstrating that continuous anodal tDCS reliably enhances cortical excitability during wakefulness ^38^.

Mechanistically, the elevated cortical excitability induced by anodal so-tDCS may have contributed to improve SO-spindle coupling and enhanced spindle activity during SO up-states. These effects likely reflect increased synaptic responsiveness, consistent with the "excitable Up-state regime" of NREM sleep proposed by Levenstein et al. ^39^. Using combined in vivo recordings and computational modeling, they demonstrated that the neocortex during NREM sleep is predominantly in an active Up-state, which is characterized by sustained neuronal firing and only transient Down-states. These findings challenged earlier conceptions of NREM sleep as a period of globally reduced neural activity and responsiveness ^40^, suggesting instead a dynamic, high-excitability cortical network in which SO down-states can arise from minor fluctuations in cortical activity. Therefore, by promoting this excitable Up-state regime during NREM sleep, and potentially amplifying rhythmic fluctuations via the oscillatory nature of the stimulation, anodal so-tDCS may have improved SO-spindle coupling.

The induced rise in cortical excitability, nevertheless, raises important questions regarding its interaction with sleep homeostasis. Sleep was shown to downscale synaptic weights and normalize excitability accumulated during wakefulness ^41^. Externally increasing excitability during NREM sleep could, in theory, interfere with this restorative process. Although we did not assess E/I balance before and after sleep, compensatory mechanisms during subsequent phases, such as REM sleep, may help restore network homeostasis. Indeed, recent evidence suggests that REM sleep can recalibrate cortical excitability and supports neural homeostasis ^27^. Future studies should determine whether anodal so-tDCS-induced excitability increase during NREM sleep is balanced by a compensatory response in later sleep stages.

Taken together, polarity of so-tDCS was a critical determinant of its functional efficacy, with anodal so-tDCS demonstrating superior impact on sleep microstructure relevant to memory.

### Chronotype modulates spindle response to anodal so-tDCS

Impact of so-tDCS on oscillatory activity has varied across individuals ^5^ and studies ^42^, likely due to differences in methodology and participant characteristics. Recent findings indicated that individual’s circadian preference for morning or evening activity influences baseline cortical excitability and the responsiveness to continuous tDCS-induced neuroplasticity during wakefulness ^43^. We therefore assessed chronotype as a potential moderator for so-tDCS during sleep and found that it strongly influenced the efficacy of anodal so-tDCS in enhancing spindle activity. While cathodal so-tDCS followed the same direction, the effect size was small. Importantly, chronotype groups did not differ in any baseline sleep measures or in cortical E/I balance prior to stimulation, indicating that both morning- and intermediate- to evening-groups were comparable in sleep oscillation characteristics and cortical excitability during NREM sleep prior to stimulation onset. The differential responsiveness emerged only after so-tDCS onset, suggesting that circadian preference is linked to the brain’s capacity to undergo plastic changes induced by so-tDCS, rather than to intrinsic differences in sleep physiology.

These results align with and extend recent findings from Salehinejad et al. ^43^, who reported that individuals exhibit higher neuroplastic changes following continuous tDCS during wakefulness if stimulation was applied at their “preferred” circadian time. In our nap paradigm, anodal so-tDCS was most effective in intermediate-to moderate evening-types, possibly because our stimulation timing (early afternoon) better coincided with their optimal neuroplastic window, i.e., peak of alertness and neural responsiveness.

## Limitations

Two limitations should be acknowledged. First, the sample size (N = 22) was modest, which may limit the generalizability of our findings and the power to detect more subtle effects. While our within-subject design enhances statistical sensitivity, a larger sample would allow more nuanced analysis of subgroup effects, particularly for chronotype. The strong correlation between chronotype and spindle enhancement suggests this factor warrants further study in larger, more diverse cohorts (i.e., including more evening chronotype individuals). Second, the cathodal condition, designed to compare neurophysiological effects of cathodal so-tDCS in an amendment to the original protocol, did not include a memory task before or after the nap. This may represent a potential confound, as pre-sleep learning has been shown to enhance subsequent SO and spindle activity ^44^, potentially biasing conditions involving pre-nap learning toward stronger oscillatory responses during sleep. However, we consider it unlikely that this influenced our findings for two reasons: (i) no differences were observed in SO or spindle metrics, their coupling, or spectral slope during the baseline N2 epoch prior to stimulation onset across conditions; and (ii) effects on SO-spindle coupling and E/I balance emerged only following so-tDCS blocks and were specific to the anodal condition. Nevertheless, future studies should control for pre-nap cognitive load to minimize potential confounding effects.

## Conclusion

In conclusion, our findings demonstrate that anodal so-tDCS during NREM sleep enhances thalamocortical oscillatory mechanisms associated with memory consolidation in older adults, whereas cathodal so-tDCS, despite modulating SO morphology, fails to engage these functional pathways and offers no observable benefit. These findings further support the concept of an excitable cortical UP-state during NREM sleep, and highlight the moderating role of individual traits such as circadian preference in brain stimulation protocols during sleep, emphasizing the need for personalized stimulation protocols.

Further, from a translational standpoint, such knowledge could contribute to interventions for cognitive aging and Alzheimer’s disease, where improving sleep oscillation quality holds therapeutic promise ^5,45^.

## Availability of data and materials

The datasets of the current study are available from the corresponding authors upon reasonable request.

## Acknowledgments

We thank M. Johns, S. König, M. Voss, M. Brechtel, J. Graefe, and K. Mahajan for their vital contribution, and all participants for taking part in the study.

## Author contributions

Buse Dikici, Data curation, Formal analysis, Validation, Investigation, Project administration, Writing – original draft, Writing – review and editing; Robert Malinowski, Software, Formal analysis, Visualization, Validation, Data curation, Supervision, Writing – review and editing; Jan-Bernhard Kordaß, Software, Formal analysis, Data curation, Supervision, Writing – review and editing; Klaus Obermayer, Conceptualization, Funding acquisition, Writing – review and editing; Julia Ladenbauer, Conceptualization, Funding acquisition, Methodology, Supervision, Project administration, Formal analysis, Writing – original draft, Visualization, Writing – review and editing; Agnes Flöel, Conceptualization, Funding acquisition, Resources, Supervision, Writing – review and editing

