## Supplemental tables for "Polarity-dependent modulation of sleep oscillations and cortical excitability in aging"

**Table S1. Baseline characteristics**

| | |
|---|---|
| n (female/male) | 22 (10/12) |
| Age (y) | 66.4 ± 6.7 (55-79) |
| Education duration (y) | 14.0 ± 3.4 (4-20) * |
| D-MEQ | 60.2 ± 9.5 (37-81) |
| AVLT, learning | 54.5 ± 9.3 (35-70) |
| AVLT, delayed recall | 1.7 ± 1.9 (-2-6) |
| Digit span, forward | 6.7 ± 2.1 (2-11) |
| Digit span, backwards | 6.0 ± 1.7 (2-8) |
| TMT, part A (time to complete,s) | 33.6 ± 8.5 (18-55) |
| TMT, part B (time to complete,s) | 78.2 ± 21.8 (41-121) |
| Letter Digit Substitution | 29.5 ± 4.8 (23-41) |
| Constructional Praxis, copy | 10.5 ± 1.1 (8-12) |
| Constructional Praxis, delayed recall | 9.0 ± 4.8 (4-28) |
| Mini-Mental-State Examination | 27.5 ± 4.7 (8-30) |
| Stroop color-word test (delay for incongruent vs neutral condition) | 98.3 ± 21.0 (72-158) |
| Verbal fluency, phonematic (no. of words) | 13.9 ± 4.2 (7-25) |
| Verbal fluency, semantic (no. of words) | 22.4 ± 5.1 (11-31) |
| Multiple choice vocabulary intelligence test | 31.7 ± 2.4 (26-35) |

Data are given as mean ± SD and range (minimum to maximum).
AVLT = auditory verbal learning test.
D-MEQ = German version of Morning-Evening-Questionnaire.
*Data available for 20/22 participants due to missing values.

**Table S2: Sleep architecture**

**Sleep architecture in 1-minute post-stimulation intervals**

| | cathodal so-tDCS | | anodal so-tDCS | | sham | | | cathodal vs. sham | anodal vs. sham |
|---|---|---|---|---|---|---|---|---|---|
| | **Mean (SD)** | **MD** | **Mean (SD)** | **MD** | **Mean (SD)** | **MD** | **p** | **corr. p** | **corr. p** |
| WASO (%) | 13.28 (16.84) | 5.28 | 18.76 (19.37) | 12.66 | 8.89 (12.56) | 4.13 | **0.066#** | 0.296 | **0.062** |
| NREM 1 (%) | 32.53 (20.25) | 32.05 | 39.91 (30.66) | 39.64 | 32.77 (21.40) | 30.18 | 0.448 | | |
| NREM 2 (%) | 47.43 (21.92) | 49.04 | 44.75 (28.97) | 44.97 | 40.74 (19.30) | 44.24 | 0.608 | | |
| NREM 3 (%) | 17.52 (17.47) | 14.84 | 9.49 (19.51) | 0.00 | 20.68 (24.51) | 14.82 | 0.143# | | |
| REM (%) | 2.53 (8.17) | 0.00 | 5.85 (15.96) | 0.00 | 5.81 (16.18) | 0.00 | 0.495# | | |
| **Sleep architecture of entire nap excluding so-tDCS/sham intervals** | | | | | | | | | |
| Total Sleep Time (min) | 43.30 (12.10) | 40.75 | 39.77 (14.17) | 38.50 | 45.07 (16.01) | 50.25 | 0.345 | | |
| Sleep Period Time (min) | 59.11 (13.43) | 63.00 | 55.57 (12.62) | 54.50 | 54.77 (16.90) | 59.00 | 0.727 | | |
| Sleep onset latency (min) | 9.30 (6.61) | 6.75 | 12.82 (9.31) | 10.25 | 9.34 (10.36) | 5.25 | 0.147# | | |
| WASO (%) | 25.74 (17.15) | 31.20 | 28.29 (18.53) | 29.35 | 17.04 (14.61) | 13.16 | 0.170# | | |
| NREM 1 (%) | 36.47 (15.21) | 37.01 | 8.36 (23.60) | 51.35 | 40.70 (20.75) | 41.79 | **0.074** | 0.459 | 0.382 |
| NREM 2 (%) | 52.59 (18.10) | 60.53 | 43.69 (20.31) | 47.10 | 42.28 (16.94) | 42.37 | 0.108# | | |
| NREM 3 (%) | 9.40 (11.80) | 5.00 | 4.99 (9.78) | 0.00 | 13.90 (17.03) | 9.42 | **0.036#*** | 0.601 | **0.046*** |
| REM (%) | 1.55 (5.11) | 0.00 | 2.97 (8.22) | 0.00 | 3.13 (10.82) | 0.00 | 0.396# | | |
| Sleep efficiency (%) | 60.50 (18.55) | 53.47 | 56.56 (20.53) | 54.20 | 62.55 (21.77) | 66.77 | 0.514 | | |
| Sleep latency N1 (min) | 9.71 (7.49) | 6.75 | 12.82 (9.31) | 10.25 | 9.66 (10.41) | 5.25 | 0.147# | | |
| Sleep latency N2 (min) | 14.36 (9.23) | 14.50 | 20.39 (14.14) | 16.75 | 16.46 (12.26) | 13.75 | **0.049#*** | 0.694 | 0.848 |
| Sleep latency N3 (min) | 24.88 (8.23) | 30.50 | 35.75 (19.15) | 38.25 | 37.88 (19.32) | 29.25 | 0.478 | | |

Sleep stages presented as percentage of total sleep time. MD = median, SD = standard deviation.
Sleep efficiency = total sleep time divided by time in bed; WASO (%) = wake after sleep onset (in min) divided by sleep period time.
Differences in sleep architecture were statistically compared between conditions with repeated-measures ANOVA, followed by planed contrasts (cathodal vs. sham; anodal vs. sham) in case of significant difference, followed by Bonferroni-Holm correction for family-wise error. Significant effects and trends towards significance are marked bold.
Please note: Stimulation/sham time periods are not included in total sleep time, sleep period time, and sleep onset latency.
### Friedman test followed by Wilcoxon signed-rank test due to skewed distribution.
*p < 0.05

**Table S3. Number of so-tDCS/sham blocks**

| Stimulation/sham block counts during the nap | cathodal so-tDCS | anodal so-tDCS | sham |
|---|---|---|---|
| 7 blocks | 2 | 3 | 3 |
| 8 blocks | 4 | 0 | 1 |
| 9 blocks | 2 | 1 | 2 |
| 10 blocks | 1 | 4 | 1 |
| 11 blocks | 3 | 4 | 2 |
| 12 blocks | 1 | 3 | 1 |
| 13 blocks | 3 | 0 | 4 |
| 14 blocks | 2 | 0 | 0 |
| 15 blocks | 4 | 7 | 8 |

Number of so-tDCS/sham blocks was determined by the participant‘s sleep in each condition.
